# The HDL particle composition determines its anti-tumor activity in pancreatic cancer

**DOI:** 10.1101/2021.07.16.452627

**Authors:** Raimund Bauer, Kristina Kührer, Florian Udonta, Mark Wroblewski, Isabel Ben-Batalla, Ingrid Hassl, Jakob Körbelin, Clemens Röhrl, Matthias Unseld, Matti Jauhiainen, Markus Hengstschläger, Sonja Loges, Herbert Stangl

## Abstract

Despite significant efforts in the last years to improve therapeutic options, pancreatic cancer remains a fatal disease and is expected to become the second leading cause of cancer-related deaths in the next decade. Late diagnosis and a complex, fibrotic tumor microenvironment produces a therapeutically hardly approachable situation with rapidly emerging resistance mechanisms. In response to this hostile microenvironment, previous research identified lipid metabolic pathways to be highly enriched in pancreatic ductal adenocarcinoma (PDAC) cells. Thereby, cholesterol uptake and synthesis was shown to promote a growth advantage to, and chemotherapy resistance for PDAC tumor cells. Here, we demonstrate that efficient, net-cholesterol removal from cancer cells, driven by high-density lipoprotein (HDL) mediated efflux, results in a significant PDAC cell growth reduction, apoptosis and a decreased PDAC tumor development *in vivo*. This effect is driven by an HDL particle composition-dependent interaction with SR-B1 and ABCA1 on cancer cells, two major lipid flux receptors, which differentially regulate cholesterol transport at the plasma membrane. Eventually, we show that pancreatic cancer patients display reduced plasma levels of HDL-cholesterol, directly translating into a reduced cholesterol efflux capacity of patient-derived plasma samples. We conclude that cholesterol depletion from PDAC cells, together with possible interventions that shunt the import and endogenous synthesis pathways of cholesterol, might represent a promising strategy to increase and complement the currently available treatment options to improve the prognosis of patients suffering from PDAC.

## Introduction

In contrast to the well-studied role of high-density lipoproteins (HDL) in cardiovascular research, its functional impact on cancer biology is less clear defined. Clinical investigations elaborated on the association of plasma apolipoprotein A1 (APOA1) / HDL levels and the risk of developing cancer, whereby the large majority of the studies reported an inverse association. For example, in randomized controlled trials of lipid-altering interventions, a significant inverse correlation between HDL cholesterol (HDL-C) and cancer incidence was found (1). Moreover, within the European Prospective Investigation into Cancer and Nutrition, the concentrations of HDL and APOA1 were inversely associated with the risk of colon cancer (2). In agreement with clinical data, preclinical studies that explored the mechanistic role of HDL in carcinogenesis predominantly attributed tumor protective functions for these lipoprotein particles. For example, B16F10 melanoma-bearing mice expressing a human APOA1 transgene exhibited reduced tumor burden, decreased tumor-associated angiogenesis, lower metastatic potential and enhanced survival. These effects were reproduced by the injection of plasma-purified human APOA1 protein into *ApoA1* knockout (KO) mice (3). To examine a causal role of reduced APOA1 / HDL levels in patients suffering from ovarian cancer, mouse *in vivo* studies with ID8 ovarian adenocarcinoma cells revealed a significant anti-tumor capacity of the human *Apoa1* transgene as well as the therapeutic administration of APOA1 mimetic peptides (4). Here, APOA1 and APOA1 mimetic peptides directly reduced the viability and proliferation of ID8 tumor cells and *cis*-platinum-resistant human ovarian cancer cell lines by the binding and removal of the mitogenic lipid lysophosphatidic acid (4).

Interestingly, a work by Cedo et al. challenged the anti-tumor activity of mature, APOA1 containing HDL. By using a model of inherited breast cancer, transgenic overexpression of human APOA1 did not result in inhibition of tumor growth. In contrast, the APOA1 mimetic peptide D-4F significantly increased tumor latency and reduced tumor outgrowth (5). Of note, APOA1 mimetic peptides as well as discoidal reconstituted HDL, which mimic pre-ß HDL particles in the circulation, are highly efficient acceptors of ATP binding cassette subfamily A member 1 (ABCA1) - mediated cholesterol efflux (6–8). In contrast, spherical, lipid rich mature HDL particles serve as high affinity ligands and donors for bidirectional, SR-B1 mediated cholesterol transport at the plasma membrane (9).

Pancreatic cancer is one of the deadliest and least therapeutically approachable malignancies with a median 5-year survival rate of approximately 5-10% (10,11). The prime reasons for this dismal prognosis are difficulties in early diagnosis and a highly diverse and hostile tumor microenvironment (TME) that promotes therapy unresponsiveness and fast developing resistance mechanisms (10,12). The TME is composed of immune cells, cancer associated fibroblasts and a dense meshwork of extracellular matrix proteins, which cause desmoplasia and a high interstitial fluid pressure (10,12). This hostile TME forces tumor cells to metabolically adapt to meet their specific requirements for proliferation, migration and invasion. Transcriptomic analyses revealed that lipid metabolic pathways are enriched in pancreatic ductal adenocarcinoma (PDAC) compared with normal pancreas. In particular, cholesterol uptake processes such as low-density lipoprotein receptor (LDLR) expression are highly activated in the malignant tissue (13). The inhibition of cholesterol uptake by PDAC cells in turn was shown to reduce cancer cell proliferation and sensitizes these cells towards chemotherapeutic interventions thereby identifying this metabolic axis as an interesting novel target for therapeutic interventions (13). Interestingly, APOA1, the main structural component of HDL and well known for its capacity to remove excess cholesterol from peripheral cells, has been identified as a potential biomarker in the detection of PDAC by comparative serum protein expression profiling (14). Although a causative link between cholesterol depletion and a reduction in pancreatic cancer malignancy seems likely, whether and how HDL particles exert an anti-tumorigenic effect in PDAC remains largely unknown.

By combining *in vitro* analyses that elucidate the impact of HDL particles on tumor cells with *in vivo* experiments using *Apoa1* knockout mice, we show that the HDL particle composition likely determines its anti-tumor activity. Reconstituted, small discoidal HDL particles displayed an increased ability to block tumor cell growth compared to lipid rich, cholesterol-laden spherical HDLs. This anti-tumorigenic capacity of small rHDL particles correlated with a lower affinity to scavenger receptor class B type 1 (SR-B1) - mediated lipid influx and a higher affinity to ABCA1-mediated cholesterol efflux. This study provides evidence for a particle composition-based anti-tumor activity of HDL in PDAC, which is at least in part regulated by an efficient cholesterol acceptor function of small, lipid poor HDLs. Therefore, we speculate that HDL-mediated cholesterol removal in combination with the blockade of cholesterol uptake from cancer cells might represent a novel, powerful mechanism to increase the efficacy of the current available concepts in PDAC treatment.

## Materials and Methods

### Animals

C57Bl6/J *Apoa1* knockout (*Apoa1* KO) mice were from The Jackson Laboratory (B6.129P2-*Apoa1*^*tm1Unc*^/J). *Apoa1* KO mice were bred heterozygously and the wildtype littermates were used as controls. All animal experiments were carried out in concordance with the institutional guidelines for the welfare of animals and were approved by the local licensing authority Hamburg (project number G36/13 and G126/15). Housing, breeding and experiments were performed with animals between 10 and 16 weeks of age under a 12h light – 12 h dark cycle and standard laboratory conditions (22 ± 1 °C, 55% humidity, food and water ad libitum).

### Cell lines and culture conditions

The murine pancreatic adenocarcinoma cells lines Panc02 and 6066 were a kind gift of Dr. Lars Ivo Partecke (Schleswig) and were maintained in RPMI medium supplemented with 10% fetal calf serum (FCS), 2 mM L-glutamine, 100 U/ml penicillin and 100 μg/ml streptomycin (complete RPMI). The human pancreatic adenocarcinoma cell line BxPC3 was from ATCC and maintained in complete RPMI medium. The murine melanoma cell line B16F10, lewis lung carcinoma cells (LLC) and the breast adenocarcinoma cell line E0771 were a kind gift of Prof. Dr. Peter Carmeliet and Prof. Massimiliano Mazzone (VIB Vesalius Research Center, KU Leuven) and maintained in DMEM medium supplemented with 10% FCS, 2 mM L-glutamine, 100 U/ml penicillin and 100 μg/ml streptomycin (complete DMEM). Cells were maintained at 37°C and 5 % CO_2_ in a humidified atmosphere and routinely tested to be mycoplasma negative (MycoAlert Mycoplasma Detection Kit, Lonza). Cells were cultured no longer than 15 passages before experimental use.

### Cholesterol depletion assays

Panc02, 6066 and BxPC3 cells (1×10^4^ cells per well of a 96 well plate) were seeded in RPMI medium containing either 2%FCS, 2% lipoprotein deficient FCS (LPDS) or 2% LPDS containing 5μM lovastatin and 100μM mevalonate for cholesterol depletion (15,16). Cells were grown for 96 h and cellular viability was determined at indicated time points using the cell proliferation reagent WST1 (Roche) according to the manufacturer’s instructions.

### Preparation of reconstituted HDL (rHDL) particles

Native human HDL was isolated from healthy donors using serial density ultracentrifugation as described (17,18). Reconstituted HDL particles were prepared according to the method of Jonas et al. (19,20). Briefly, 1mg of native HDL was delipidated twice using 5mL ethanol:diethyl ether (3:2). The supernatant was discarded and the remaining solvents were evaporated with nitrogen gas. Phosphatidylcholine (PC), cholesterol (C) and cholesteryl-palmitate (CE, in chloroform:methanol (2:1)) were combined in specific molar ratios and the solvents were evaporated with nitrogen gas. The dried lipids were resuspended in 200 μL buffer A (150mM NaCl, 0.01% EDTA, 10mM Tris/HCl, pH 8.0) and 50 μL of a 30 mg/mL sodium deoxycholate solution was added to disperse the lipids. The mixture was stirred at 4°C for two hours. Delipidated HDL was dissolved in 250 μL of buffer A. Both suspensions were mixed in a glass vial and stirred at 4°C overnight. The suspension was then filtered twice through a 4 mL 3K Amicon filter tube and the protein concentration was determined. Reconstituted HDL was overlaid with nitrogen gas, and stored at 4°C. To prepare tracer-labeled rHDL, 25 μCi ^3^H Cholesteryl oleate (Perkin Elmer, NET746L001MC) and 12.5 μCi ^14^C Cholesterol (Amersham, CFA128) dissolved in toluene were added to the lipid mixture (for an equivalent of delipidated APOA1 of 500μg) before evaporation.

### (r)HDL treatments of pancreatic adenocarcinoma cell lines

Panc02 and 6066 cells were starved in a T75 flask overnight in RPMI medium containing 0.1% FCS. Next, 1×10^4^ cells were seeded per well of a 96 well plate in RPMI medium containing 2% FCS with the addition of indicated HDL particles (75 μg/ml). To inhibit SR-B1, BLT1 (0.5μM) was added 30 min before the addition of HDL to the cell suspension. For ABCA1 activation, TO901317 (or DMSO control) was added at the indicated concentrations directly to the starvation medium the day before the assay and throughout the experiment.

### Quantitative PCR

Total RNA was isolated from cells or murine liver tissue using the Relia Prep RNA Tissue Miniprep System (Promega) according to the manufacturer’s instructions. One μg of RNA was reverse transcribed into single stranded cDNA (Go Script Reverse Transcription System, Promega) and subsequently used for qPCR analyses on a Step One Plus real time PCR detection system (Applied Biosystems). Expression levels of genes of interest were normalized to hypoxanthine guanine phosphoribosyl transferase (*Hprt*) and the relative fold gene expression compared to control was calculated using the 2^−(ΔΔCt)^ method.

### Western blotting

0.5 μl of mouse plasma or 20 μg of Panc02 RIPA total protein extracts were analyzed by reducing, SDS-PAGE (8% PAA gel) and transferred to nitrocellulose membranes. Nonspecific binding sites were blocked with TBS (20 mM Tris-HCl, pH 7.4, 137 mM NaCl) containing 5% (w/v) fatty acid-free BSA and 0.1% Tween-20 (blocking buffer) for 1 h at room temperature. Proteins of interest were detected with antibodies for APOA1 (in-house produced rabbit polyclonal anti-human APOA1 antibody), ABCA1 (MAB10005, Merck) and β-ACTIN (clone AC-74, Sigma Aldrich) followed by incubation with HRP-conjugated secondary antibodies and development with the enhanced chemiluminescence protocol (Pierce).

### Cholesterol flux assays

To measure cholesterol efflux from Panc02 or 6066 cells to HDL particles, 0.1×10^6^ cells were seeded in 900 μl of complete RPMI per well of a 12 well plate and incubated for 24 h at 37°C and 5 % CO_2_. Next, cells were trace-labeled for another 24 h with ^3^H cholesterol (Perkin Elmer, NET139) by adding 100 μl of complete RPMI containing 5μCi ^3^H cholesterol / ml per well. The next day, cells were washed twice with warm RPMI medium containing 0.1% FCS and once with 1 ml of warm PBS. After carefully removing the medium, 500 μl of 0.1% FCS-containing RPMI containing HDL particles (10 μg/ml) to the cells. When analyzing the cholesterol acceptor capacity of human plasma samples, instead of HDL particles, 10 μl (2%) of plasma was added to 500 μl of 0.1% FCS-containing RPMI. In SR-B1 inhibition studies, BLT1 (1μM) was added 1 h prior to the addition of efflux acceptors. For the activation of ABCA1 expression, the LXR agonist TO901317 (5μM) was added to cells 48 h prior to the addition of efflux acceptors. Efflux acceptors were incubated for 8 h with the cells. To analyze the transfer of cholesterol and its esters from HDL to the cellular compartment, tracer-labeled HDL particles (10 μg/ml) were again diluted in 0.1% FCS containing RPMI medium and incubated for 8 h with the cells. Supernatants are collected and cleared from cellular debris by centrifugation at 10.000 x g for 10 min at room temperature. Cells were washed twice with PBS and lysed by the addition of 500 μl 0.1N NaOH. 200 μl of either supernatant or cell lysate were mixed with 8 ml of Ultima Gold scintillation cocktail and analyzed by scintillation counting.

### Adeno-associated viral (AAV) particle production

The production of liver-targeting AAV particles of the serotype AAV2.8 was performed as previously described in detail (21-23). Briefly, the full length murine APOA1 cDNA was inserted into a pAAV-MCS plasmid containing AAV inverted terminal repeats (ITR) using BstBI (fwd) and BsrGI (rev) restriction enzyme sites. Together with a pAAV rep2 cap8 transfer plasmid and an AdpXX6 helper plasmid, HEK cells were co-transfected and virus particles purified from cell pellets and supernatants using iodixanol density gradients (21,23).

### *In vivo* experiments

To compare tumor growth kinetics in WT and *Apoa1* KO mice, 0.5 × 10^6^ Panc02, B16F10 or LLC cells were implanted subcutaneously into the right flank of mice. E0771 cells were implanted orthotopically into the second mammary fat pad. Tumor size was measured with a digital caliper and the volume was calculated using the formula V= (length^2^ × width) / 2. For histological analyses, pimonidazole (1 mg, i.p.) was injected 2 h before sacrifice. For AAV-mediated reconstitution of hepatic APOA1 levels in *Apoa1* KO mice, 2×10^11^ AAV2.8 particles encoding the full length murine APOA1 mRNA (AAV-APOA1) were administered intravenously 5 days prior to tumor cell inoculation. For rHDL injection studies, tumors were grown to a size of 100 mm^3^. Afterwards, mice received intravenous injections of either 0.2 mg rHDL (PC:C:CE:APOA1 = 100:12.5:0:1; ZLB Behring, a kind gift of Prof. Matti Jauhiainen) or an equivalent volume of sterile PBS every 72 h.

### Collection of plasma samples and analysis of lipid parameters

Murine blood samples were obtained by retroorbital bleeding and collection of blood into precoated EDTA tubes (Sarstedt). Human blood samples were collected into precoated EDTA tubes from pancreatic cancer patients under informed consent and strict adherence to institutional guidelines of the Medical University of Vienna (Ethik Votum 1035/2020). Blood samples were immediately centrifuged for 15 min at 3000 × g at room temperature and plasma samples were collected and frozen at −80°C until further use. Total plasma triglycerides, cholesterol and HDL-C were measured with the Triglyceride FS, Cholesterol FS and HDL-C Immuno FS kit systems (DiaSys).

### Analysis of intratumoral hypoxia

Tumor samples were fixed overnight in 4% paraformaldehyde at 4 °C and embedded in paraffin. Paraffin sections (4 μm) were stained with antibodies to detect tumor hypoxia (pimonidazole, HP3-1000kit) as previously described (24). For morphometric analysis, 8-10 optical fields per tumor section were acquired using a Zeiss Axio Scope A1 and images were analyzed using the NIH Image J software.

### Flow cytometry

Flow cytometric analysis of enzymatically digested tumor tissue was essentially performed as described in (24). MRC1+ macrophages were gated as PE-Cy7 CD11b^+^ (clone M1/70 BD Bioscience), PE F4/80^+^ (clone BM8, Biolegend) and FITC CD206 (MRC1)^+^ (clone C068C2, Biolegend). Granulocytic myeloid derived suppressor cells (GMDSC) were gated as PE-Cy7 CD11b^+^, PE Ly6-G^+^ (clone 1A8 BD Bioscience) and PerCP-Cy5.5 Ly6-C^int^ (clone Hk1.4, eBioscience). T cells were characterized as APC CD3^+^ (clone 17A2, eBioscience) and either eFluor 450 CD8a^−^(clone 53-6.7, eBioscience) FITC CD4^+^ (clone GK1.5, Biolegend) for T helper cells or vice versa for cytotoxic T cells. DAPI was used as a viability stain.

### Detection of cellular apoptosis

Trypsinized and washed Panc02 cells were stained with the FITC Annexin V Apoptosis Detection Kit with 7-AAD (Biolegend) and analyzed using the CytoflexS Flow Cytometer (Beckman Coulter).

### Statistics

Data represent mean ± SEM of representative experiments. To compare the means of two groups, an unpaired, two-tailed student’s t-test was used. Multiple comparison testing in experiments with more than two groups was performed using one-way ANOVA unless otherwise stated. Statistical significance was assumed when p<0.05.

## Results

### Cholesterol depletion and small, discoidal HDL particles efficiently inhibit the growth of PDAC cell lines

Previous studies indicate that cancer cells show increased sensitivity towards cholesterol depletion due to their high need of cholesterol for cellular growth (25). In pancreatic cancer, the blockade of cholesterol uptake and the depletion of cholesterol availability via statins have been shown to reduce pancreatic cancer risk in preclinical as well as clinical settings (13,26). In accordance, reducing cholesterol availability by culture of cells in lipoprotein deficient serum (2% LPDS) decreased the viability of murine pancreatic adenocarcinoma cells Panc02 (Figure 1A). Cholesterol depletion in Panc02 cells by lovastatin further reduced cellular viability (Figure 1A). By comparing murine pancreatic cancer cell lines Panc02, 6066 (27,28) and the human pancreatic cancer cell line BxPC3 regarding their sensitivity towards cholesterol depletion, all three cell lines demonstrated reduced cellular viability, with Panc02 cells being the most sensitive (Figure 1B). HDL particles serve as important acceptors of cellular cholesterol, with the capacity to remove excess cholesterol from peripheral cells (29). Importantly, and within the HDL pool, cholesterol efflux capacity differs according to the HDL particle size, lipid and protein composition and the specific affinities for cellular efflux receptors such as ABCA1, ABCG1 or SR-B1 (9,30,31). By comparing the impact of native human spherical HDL (predominantly cholesterol donors) with small, lipid-poor reconstituted HDLs (rHDL, predominantly cholesterol acceptors) on the proliferative capacity of Panc02 cancer cells, we observed that under serum starvation conditions, rHDLs reduced cellular viability more efficiently (Figure 1C). Although HDLs decreased Panc02 viability also with increasing concentrations of serum in the cell culture medium, the particle-specific effect was attenuated (Figure 1C).

**Figure 1.**
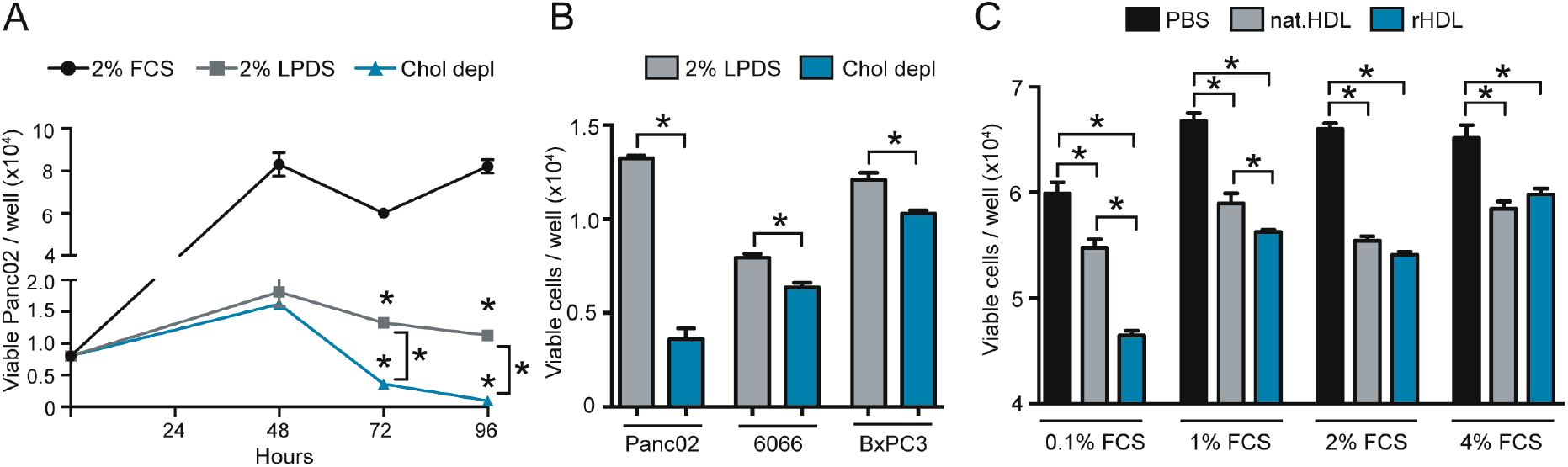
Pancreatic cancer cells are sensitive towards cholesterol depletion and rHDL particles. **A**, Panc02 cells were grown in RPMI medium containing either 2% FCS, 2% LPDS or 2% LPDS with the addition of 5μM lovastatin and 100μM mevalonate. Viability of cells was determined every 24 h using a WST1 viability assay (n=4/4/4; *p<0.05; one-way ANOVA). **B**, Viability was determined of Panc02, 6066 and BxPC3 pancreatic cancer cells after culture of cells for 72 hours in either RPMI medium with 2% LPDS or cholesterol depleted medium (n=4/4; *p<0.05; unpaired t-test). **C**, Panc02 cells cultured in RPMI medium with increasing concentrations of FCS were treated for 48 h with PBS, HDL isolated from human plasma of a healthy donor (75μg/ml) or reconstituted HDL (75μg/ml, molar ratio of PC:C:CE:APOA1 of 100:12.5:0:1, ZLB Behring) and viability was determined using a WST1 assay (n=4/4/4; *p<0.05; one-way ANOVA).

### The HDL particle composition affects viability and apoptosis of PDAC cells

To evaluate a potential particle-specific anti-tumor effect of HDL, we produced rHDL particles with varying lipid compositions to either mimic small, discoidal HDL particles (rHDL1) or spherical, lipid-rich HDLs (rHDL2; Figure 2A). Treatment of Panc02 and 6066 murine PDAC cell lines revealed that small HDL discs (rHDL1) induced a profound reduction in cellular viability, whereas treatment with nHDL and rHDL2 resulted in only a mild attenuation in pancreatic cancer cell viability (Figure 2B and C). As persistent cholesterol starvation can induce cellular apoptosis (32,33), we analyzed apoptosis rates in HDL-treated pancreatic cancer cells (representative FACS blots are shown in Figure 2D). Whereas all HDL particles promoted a slight decrease in the frequency of living cells, only rHDL1 treatment led to a significant reduction of the live cell population (Figure 2E). Additionally, HDL treatment increased early apoptosis rates compared to control irrespective of the particle composition (Figure 2F). In agreement with the data from viability assays (Figure 2B and C), rHDL1 was able to significantly expand the late apoptotic cell pool, indicating substantial cancer cell killing activity of this small, lipid-poor HDL particle (Figure 2G).

**Figure 2.**
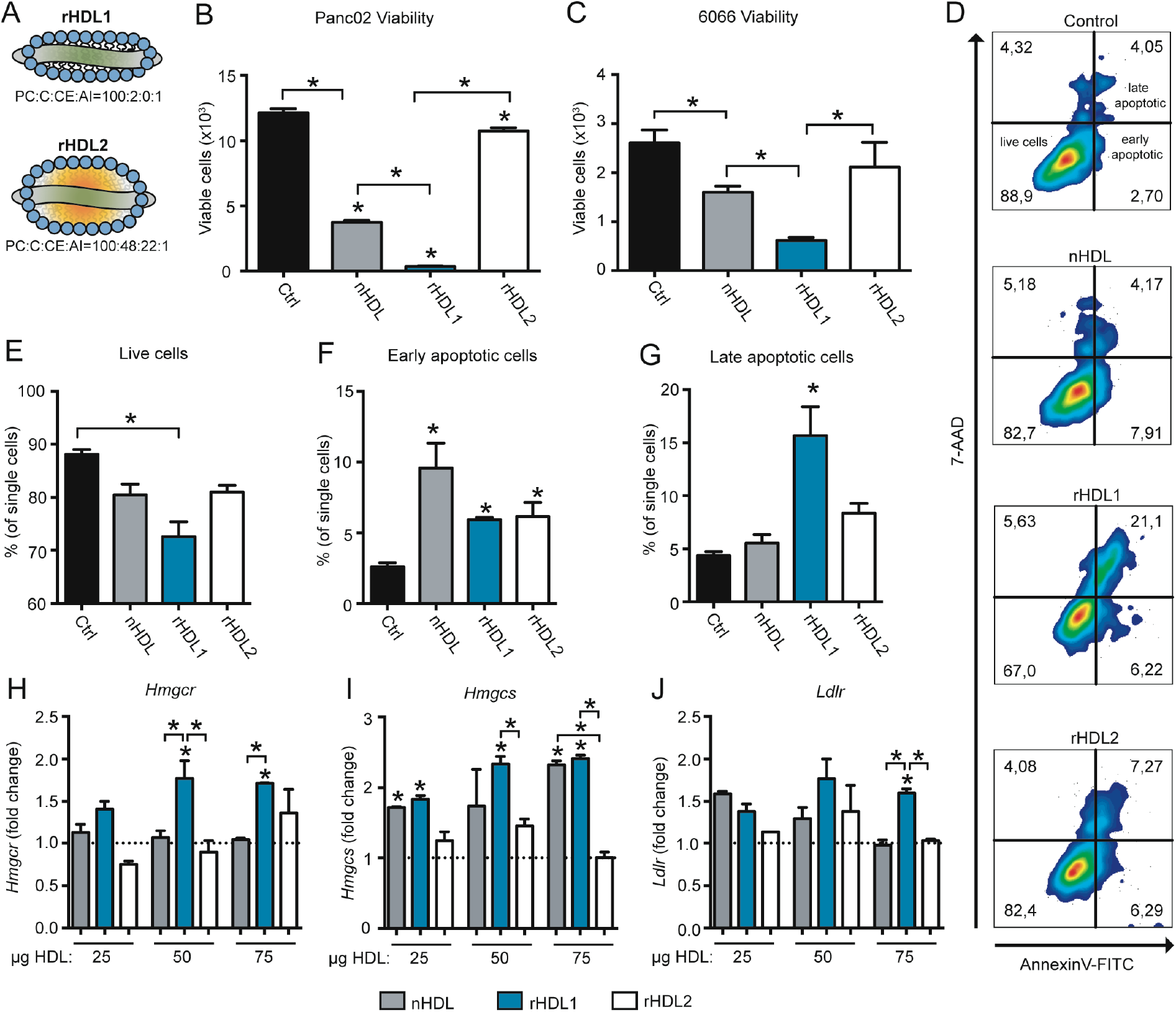
The HDL particle composition differentially affects pancreatic cancer cell growth characteristics. **A,** HDL particles were reconstituted using the indicated molar ratios of phosphatidylcholine (PC), cholesterol (C), cholesterol ester (CE) and apolipoprotein A-I (APOA1). **B, C**, Following a 24 h starvation period, Panc02 and 6066 cells were cultured in RPMI medium with 2% FCS in the presence of either PBS, native human HDL (nHDL), or the indicated rHDL particles (75μg/ml) for 48 h. Afterwards, viability was determined using a WST1 assay (n=4 replicates per group; *p<0.05; unpaired t-test). **D**, Representative flow cytometry blots of Panc02 cells treated with different HDL particles (75μg/ml) stained for the detection of apoptosis using 7-AAD and Annexin-V. **E-G**, quantification of live-, early apoptotic-, and late apoptotic cells from flow cytometry experiments shown in **D** (n=3 replicates per group, *p<0.05; one-way ANOVA). **H-J**, Panc02 cells were starved for 8 h and afterwards treated with indicated concentrations of different HDL particles for 16 h. qPCR experiments show relative mRNA levels of *Hmgcr*, *Hmgcs* and the *Ldlr* (n=3 replicates per group; *p<0.05; one-way ANOVA).

The depletion of cellular cholesterol pools activates endogenous cholesterol synthesis pathways as well as LDLR expression for the exogenous uptake of cholesterol (34,35). Therefore, we hypothesized that rHDL treatment might increase transcription of key enzymes of the cholesterol synthesis / uptake machinery. Indeed, and in contrast to control-, nHDL- and rHDL2-treated cells, rHDL1 particles significantly induced the expression of 3-hydroxy-3methyl-glutaryl-coenzyme A reductase (*Hmgcr*) at 50 and 75 μg/ml (Figure 2H). While rHDL1 and nHDL dose-dependently increased mRNA levels of the hydroxymethylglutaryl-CoA synthase (*Hmgcs*), rHDL2 failed to do so (Figure 2I). Furthermore, the lipid-poor rHDL1 particles increased LDLR gene expression at 50 and 75 μg/ml. Together, these data indicate that rHDL1 particles reduce viability and induce apoptosis of pancreatic cancer cells, paralleled by an induction of the cellular cholesterol synthesis- and import machinery.

### The HDL particle composition determines its net cholesterol efflux capacity from PDAC cells

These observed effects suggested efficient cholesterol depletion of cancer cells when treated with rHDL1 particles, likely facilitated via cholesterol efflux from cellular cholesterol pools to extracellular HDL. To measure the cholesterol efflux capacity of different HDL species, we labeled Panc02 and 6066 cells with ^3^H cholesterol and analyzed the transfer of the radiotracer onto HDL. As anticipated from previous results, rHDL1 was able to remove significantly more cholesterol from cancer cells compared to nHDL. However, the cholesterol efflux capacity of lipid-rich rHDL2 particles even exceeded the one of rHDL1 (Figure 3A). Cholesterol efflux is primarily regulated by cell surface receptors such as SR-B1 and family members of the ABC-transporters such as ABCA1 and ABCG1. While expression levels of SR-B1 are high in Panc02 cells, ABCA1 expression is rather low and ABCG1 mRNA levels are hardly detectable (Supplementary Figure 1). Therefore, and to get a more detailed view of the cholesterol transport properties of the different HDL particles, we first analyzed cholesterol efflux in the presence or absence of the SR-B1-blocking small molecule inhibitor BLT1 (36). Pretreatment of Panc02 cells with BLT1 significantly reduced cholesterol efflux to nHDL and rHDL1 particles. In contrast, efflux of ^3^H cholesterol towards rHDL2 particles was only moderately affected (Figure 3B). As SR-B1 mediates bi-directional lipid transfer (9), we synthesized rHDL particles containing both ^3^H cholesteryl-ester and ^14^C cholesterol (Figure 3C) and used those particles to analyze lipid influx by measuring the accumulation of intracellular radiotracer molecules. Although containing the same amount of radiotracer compared to rHDL1, the influx of free cholesterol was significantly increased when Panc02 cells were incubated with the lipid-laden rHDL2 particles. BLT1-mediated SR-B1 inhibition blunted the influx of free cholesterol to a similar extent (Figure 3D). Interestingly, lipid-laden rHDL2 particles substantially exceeded rHDL1 particles in their ability to transfer cholesteryl-oleate onto pancreatic cancer cells. Similar to the data observed for cholesterol efflux, BLT1 blocked this effect only to a small extent (Figure 3E). These data identify rHDL2 particles to be more efficient in mediating lipid exchange with pancreatic cancer cells compared to lipid-poor rHDL1 particles and an overall reduced efficacy of BLT1 to block rHDL2-mediated lipid flux. Of note, while rHDL1 particles again showed the highest apoptosis-inducing capacity, BLT1 treatment of Panc02 cells even further induced early and late apoptotic cells in the presence of rHDL1, but not of rHDL2 particles (Figure 3F and G). One explanation could be that compared to nHDL and rHDL1, the affinity of rHDL2 might be higher towards SR-B1, thereby reducing the ability of BLT1 to block lipid transport. This phenomenon of high affinity of lipid-rich HDLs towards SR-B1 has been previously described (reviewed in (9)), which might further be exacerbated by the high expression levels of SR-B1 in Panc02 cells.

**Figure 3.**
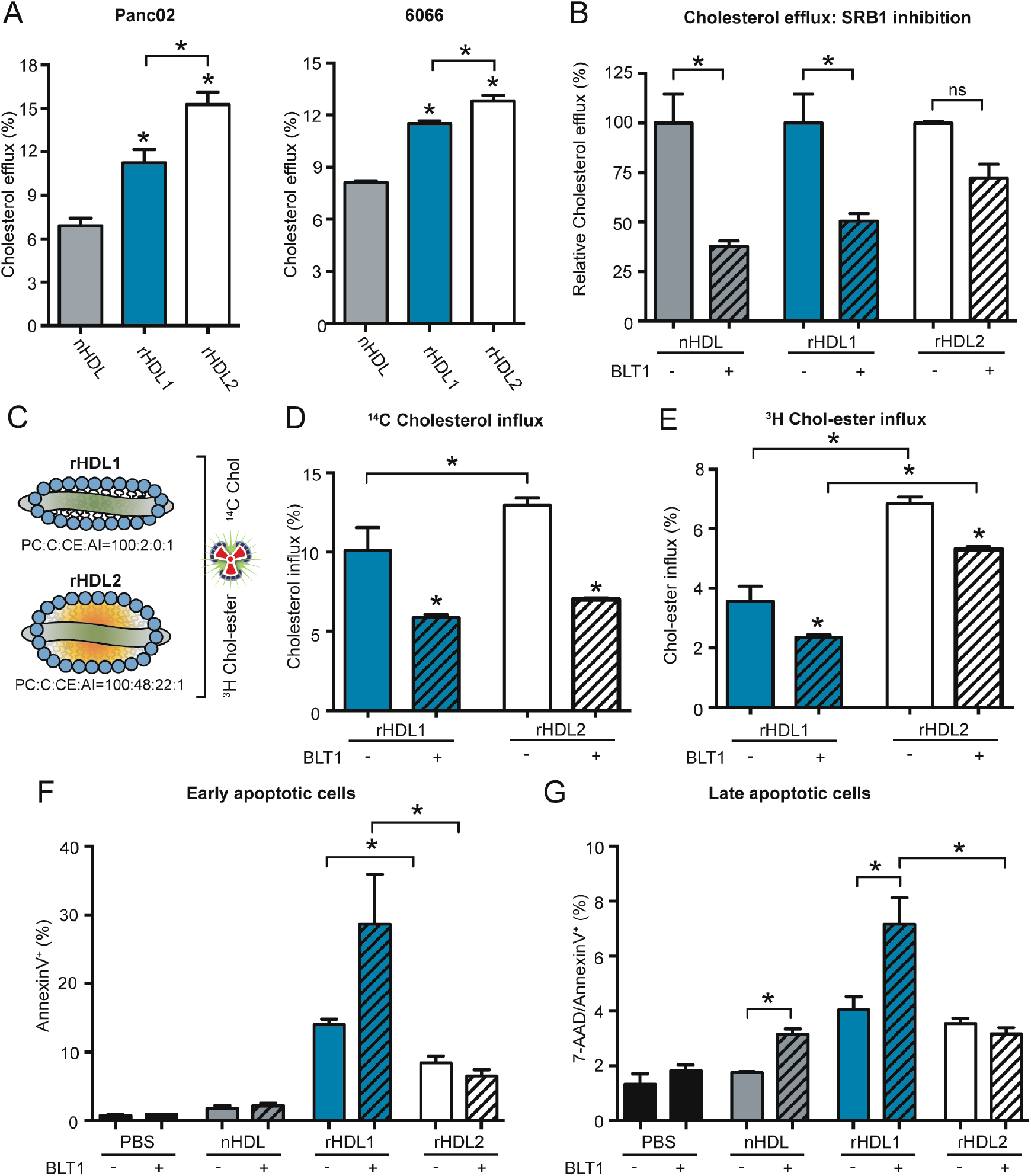
SR-B1 influences cholesterol efflux and anti-tumor properties of rHDL particles. **A,** ^3^H cholesterol-loaded Panc02 and 6066 cells were subjected to cholesterol efflux assays with indicated HDL particles (10μg/ml) for 8 h. Cholesterol efflux is shown as % of transferred tracer from cells to HDL particles compared to control conditions (n=4; *p<0.05; one-way ANOVA). **B,** relative, particle specific cholesterol efflux capacity was analyzed in the absence or the presence of BLT1 (n=4; *p<0.05; one-way ANOVA). **C,** schematic representation of tracer-labeled rHDL particles. **D**, **E**, ^14^C cholesterol and ^3^H cholesteryl ester influx, respectively, from rHDL to Panc02 cells in the absence or presence of BLT1(n=4; *p<0.05; one-way ANOVA). **F**, quantification of early and **G**, late apoptotic cells upon treatment of Panc02 cells with indicated HDL particles in the absence or presence of BLT1 (n=3; *p<0.05; unpaired t-test).

### The LXR agonist TO901317 increases rHDL1-specific cholesterol efflux and apoptosis

The rHDL1-specific increase in apoptosis upon inhibition of SR-B1 points towards the involvement of SR-B1-independent mechanisms that mediate the anti-neoplastic effect of rHDL1. As small and lipid-poor HDL particles are highly efficient substrates for ABCA1-mediated cholesterol efflux, we analyzed the potential role of ABCA1 in the cholesterol efflux-driven anti-poliferative effects of rHDL1 particles. To manipulate ABCA1 protein levels in Panc02 cells, we used the LXR-agonist TO901317 (TO, Figure 4A). As expected, TO treatment led to an increase of cholesterol efflux to rHDL1 particles of 35%. Although cholesterol efflux to rHDL2 particles was also increased in the presence of TO, this effect was significantly weaker compared to rHDL1-mediated efflux (Figure 4B). Importantly, only rHDL1 particles reduced the amount of intracellular ^3^H cholesterol in the presence of TO, pointing towards an ABCA1-centered rHDL1-specific depletion of cholesterol pools in Panc02 cells (Figure 4C). Finally, SR-B1 inhibition of TO-treated, ABCA1-expressing cells showed a profound pro-apoptotic effect on Panc02 cells in the presence of rHDL1 particles, which was blunted when rHDL2 particles were used (Figure 4D).

**Figure 4.**
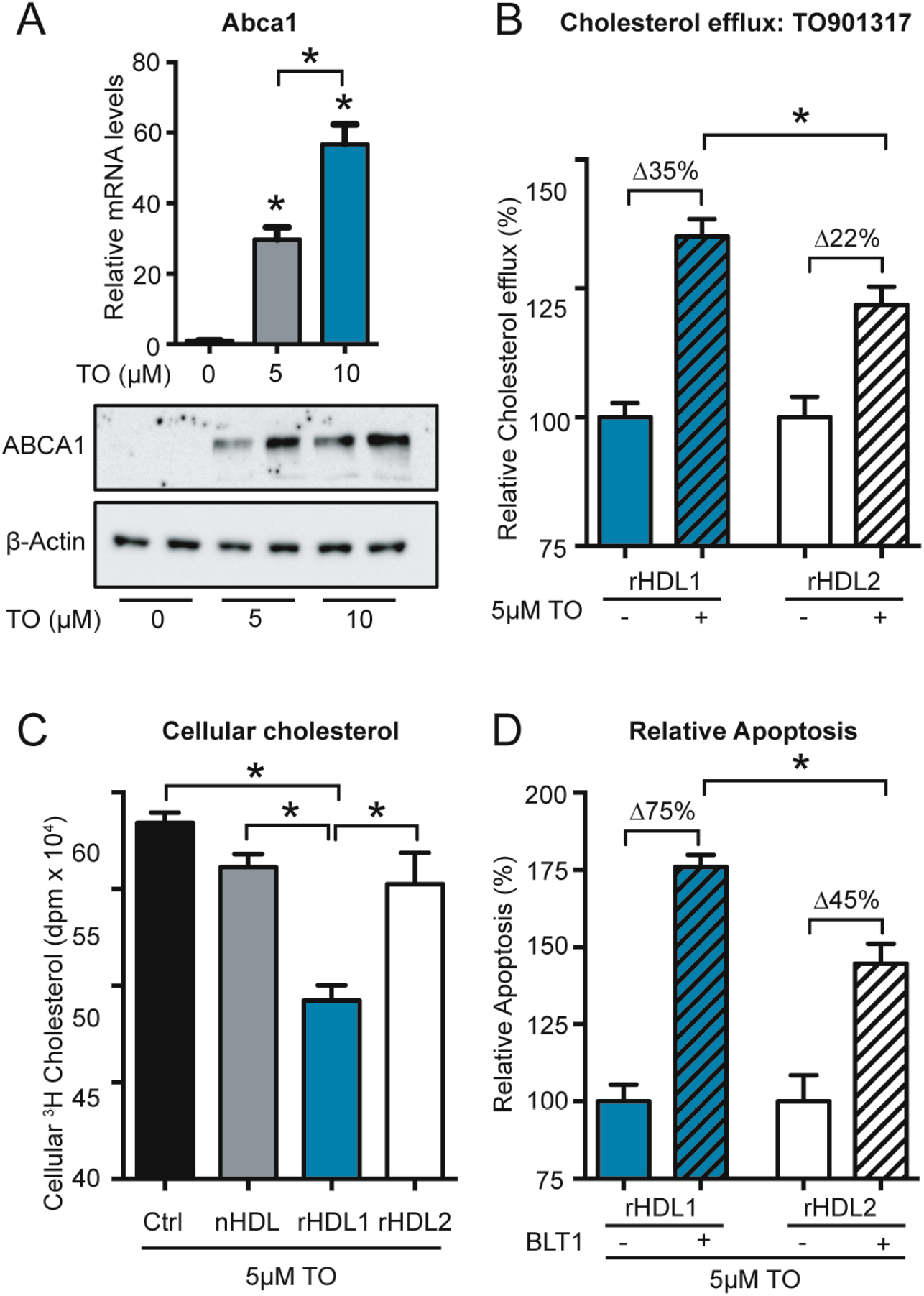
The LXR agonist TO901317 increases cholesterol efflux and apoptosis-inducing properties of small, lipid poor rHDL. **A,** TO901317 induces ABCA1 mRNA and protein levels in Panc02 cells (n=3; *p<0.05; one-way ANOVA). **B**, relative, particle specific cholesterol efflux from Panc02 cells in the absence or presence of TO901317 (n=4; *p<0.05; unpaired t-test). **C**, ^3^H cholesterol accumulation in Panc02 cells in the presence of TO901317 following efflux to indicated HDL particles (n=4; *p<0.05; one-way ANOVA). **D**, relative, particle-specific apoptosis rates of TO901317-treated Panc02 cells induced by either rHDL1 or rHDL2 in the absence or presence of BLT1 (n=3; p<0.05; unpaired t-test).

Together, lipid poor, discoidal-like rHDL1 particles induced a significant pro-apoptotic effect in Panc02 cells by unidirectional cellular cholesterol removal via ABCA1. In contrast, the pro-apoptotic effect was diminished when lipid-rich rHDL2 particles or native HDL isolated from human plasma were used. The previously reported high affinity of those particles to SR-B1 as well as their increased efficacy in mediating bidirectional lipid flux reduces the net cholesterol-removing capacity of those particles, thereby making them less effective in killing pancreatic cancer cells.

### Liver-specific AAV-mediated APOA1 expression and rHDL injections reduce tumor burden in Panc02-bearing *ApoA1* KO mice

To demonstrate this anti-tumor effect of HDL particles *in vivo*, we first compared tumor growth kinetics of WT and APOA1 deficient mice (*Apoa1* KO), which exhibit dramatically reduced plasma HDL levels (37). Of note, we were unable to detect significant differences in tumor growth and tumor weight in the Panc02 tumor model as well as in the B16F10, LLC and E0771 tumor models, which demonstrates an insignificant anti-tumor effect of mature, endogenous HDL particles at least in the murine system (Supplementary Figure 2A-D). Next, and as a consequence of the data obtained from *in vitro* experiments, we decided to artificially introduce APOA1 / rHDL particles into tumor-bearing *Apoa1* KO mice. Therefore, we expressed murine APOA1 in the liver of Panc02 tumor-bearing mice using adeno-associated viral particles (AAV-APOA1). Robust expression levels of the APOA1 mRNA were detected in in the liver of end stage Panc02 tumor-bearing WT and *Apoa1* KO AAV-APOA1 mice, whereas APOA1 mRNA levels were absent from livers of *Apoa1* KO mice (Figure 5A). Western blot analysis revealed the absence of APOA1 from plasma samples of *Apoa1* KO mice, whereas APOA1 protein was readily detectable in the plasma 5 days post AAV injection and further increased after 14 and 21 days (Figure 5B). AAV-mediated APOA1 expression significantly increased HDL-C levels compared to APOA1 deficient mice, although HDL-associated cholesterol levels clearly remained below those in WT mice (Figure 5C). Importantly, AAV-APOA1 expression in *Apoa1* KO mice significantly reduced Panc02 tumor growth kinetics as well as tumor weight at experimental end stage (Figure 5E). Finally, and to examine a therapeutic potential of rHDL particles *in vivo*, we compared Panc02 tumor growth in WT mice, *Apoa1* KO mice receiving PBS and *Apoa1* KO mice receiving intravenous injections of rHDL (0.2 mg per injection every other day). Thereby, rHDL reduced Panc02 tumor weight significantly compared with *Apoa1* KO mice, pointing towards a moderate anti-tumor effect of rHDL particles also *in vivo* (Figure 5F).

**Figure 5.**
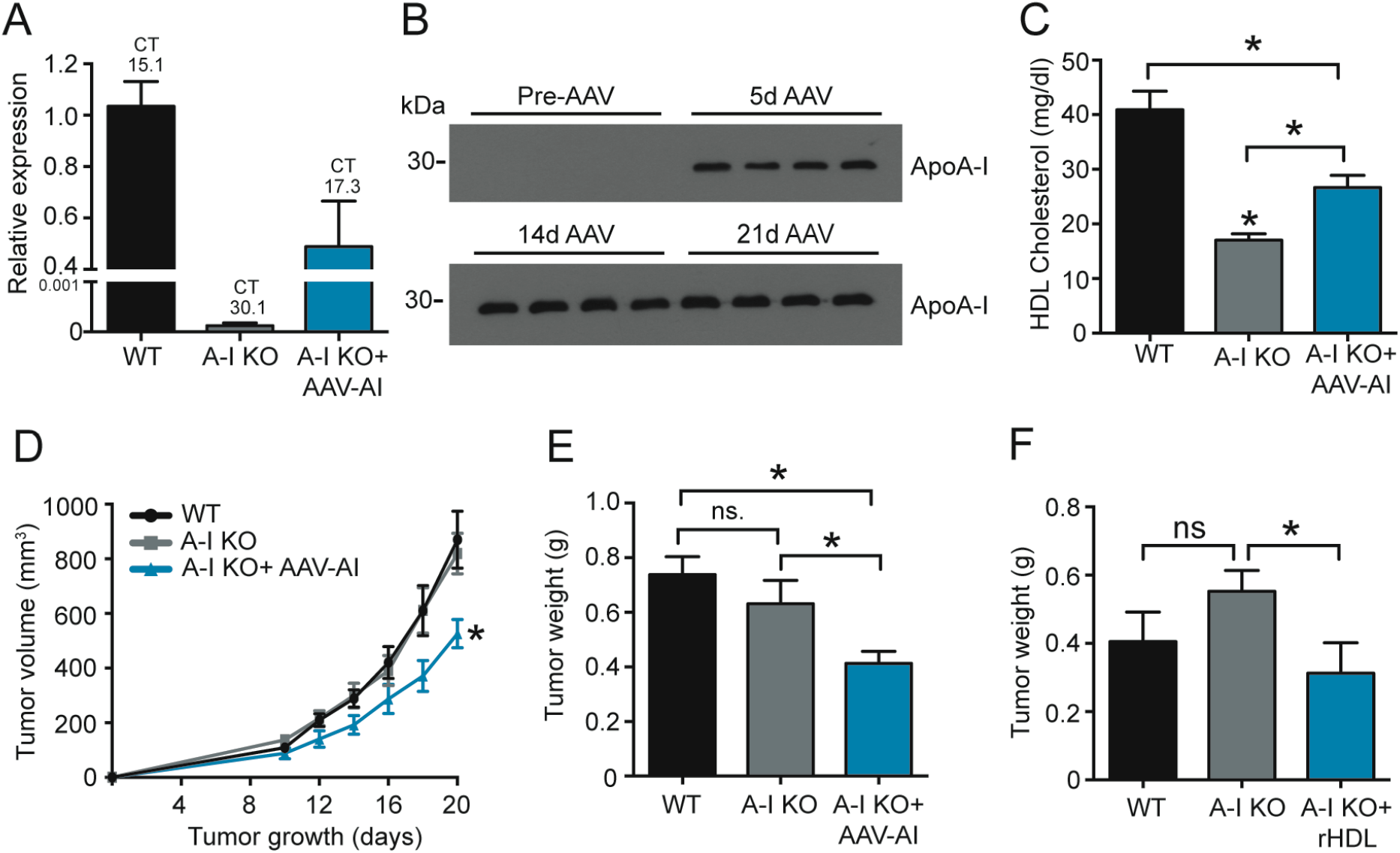
AAV-mediated APOA1 reconstitution / rHDL injection reduces Panc02 tumor growth in *Apoa1* KO mice. **A,** mRNA expression levels of APOA1 in liver tissue of Panc02 bearing WT, *Apoa1* KO and AAV-APOA1 reconstituted *Apoa1* KO mice at experimental end stage (n=8/6/7). **B,** Western blot analysis of APOA1 protein levels in plasma of *Apoa1* KO mice prior and post AAV-APOA1 injection. **C**, HDL-C levels in Panc02 tumor bearing mice at experimental end stage (n=8/5/7; *p<0.05; one-way ANOVA). **D**, Panc02 tumor growth kinetics and **E**, tumor weight at experimental end stage (n=8/5/7; *p<0.05; unpaired t-test). **F**, Panc02 tumor weight of WT, *Apoa1* KO and rHDL (0.2mg per injection; PC:C:CE:APOA1 = 100:12.5:0:1, ZLB Behring) - injected *Apoa1* KO mice at experimental end stage (n=7/7/7; *p<0.05; unpaired t-test).

As APOA1 (mimetic peptides) / HDL has been previously demonstrated to affect tumor angiogenesis as well as tumor associated immune cell populations (3), we measured intratumoral hypoxia and immune cell infiltration. Thereby, we found that AAV-mediated APOA1 expression did not affect tumor associated hypoxia, a surrogate marker for tumor angiogenesis (Supplementary figure 3A and B). Flow cytometric analysis of tumor associated immune cell populations showed an increase in MRC1^+^ M2-polarized Macrophages in the *Apoa1* KO background, which was not substantially reverted by reintroduction of APOA1 (Supplementary Figure 3C). Other tumor-associated immune cell populations such as myeloid-derived suppressor cells (MDSC), CD8^+^cytotoxic T cells and CD4^+^ T-helper cells remained unchanged among all three groups of mice (Supplementary Figure 3D, E, F). Together, we conclude that in the here applied model of PDAC, HDL neither rendered tumor-associated hypoxia / angiogenesis nor did it change infiltration of immune cell populations. Therefore, we speculate that in parallel to the data from *in vitro* experiments, the increased cholesterol efflux capacity of artificially introduced HDL particles mediates the observed anti-tumor effect.

### Decreased HDL-C and decreased efflux capacity of plasma samples from PDAC patients

To test our hypothesis in a clinical relevant setting, we collected plasma samples from pancreatic cancer patients and analyzed lipid parameters and cholesterol efflux capacities. Although pancreatic cancer plasma samples showed no difference in triglyceride and total cholesterol levels (Figure 6A and B), HDL-C levels were significantly decreased when compared to plasma samples of a cohort of healthy volunteers (Figure 6C). Of note, this drop in HDL-associated cholesterol also translated into decreased cholesterol efflux capacity of the patient plasma samples from Panc02 cells (Figure 6D). Accumulation of intracellular cholesterol can either be regulated by uptake of extracellular cholesterol via the LDLR or endogenous synthesis of cholesterol from acetyl-CoA precursor molecules. *In silico* Kaplan Meier analyses with the UCSC Xena database (38) further revealed a potential dysregulation of cholesterol homeostasis in PDAC, showing an inverse association of LDLR and HMGCS expression levels with overall survival in a cohort of pancreatic cancer patients (Figure 6E and F). This inverse correlation of the LDLR and HMGCS with overall survival persisted with high significance when analyzing the TCGA Pan-cancer database (Supplementary Figure 4A and B). In contrast, expression of HMGCR showed no association with survival data (Supplementary Figure 4C and D). Interestingly, expression levels of *MYLIP*, the gene encoding the LDLR-degrading ubiquitin E3 ligase IDOL, showed a significant, positive association with overall survival parameters in the TCGA Pan-cancer database and inversely correlated with LDLR expression levels in the analyzed primary tumor samples (Supplementary Figure 4E and F). These data further underline the importance of cholesterol availability for pancreatic tumor growth and indicate that the depletion of intracellular cholesterol pools might hold potential as a therapeutic strategy to improve the prognosis of pancreatic cancer patients.

**Figure 6.**
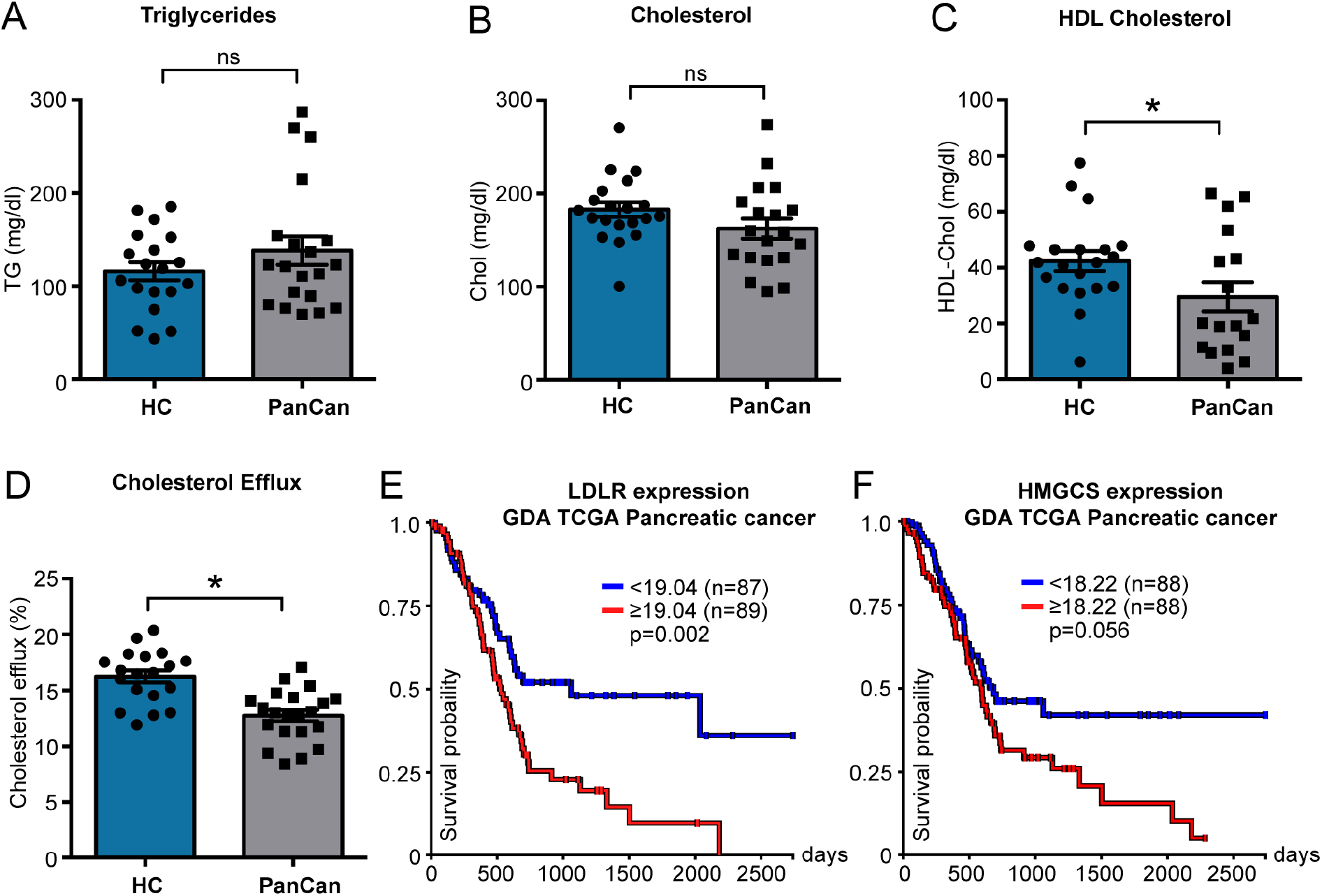
Pancreatic cancer patients show decreased plasma HDL-C and cholesterol efflux capacities compared to healthy donors as well as increased survival probability with lower tumor-associated LDLR and HMGCS expression levels. **A,** plasma total triglyceride levels, **B,** total cholesterol levels, **C**, HDL-C and **D,** plasma efflux capacity were determined in plasma samples from a cohort of healthy volunteers and pancreatic cancer patients in late stages of their disease (n=19/19; *p<0.05; unpaired t-test). **E, F,** Kaplan Meier overall survival plots for LDLR and HMGCS expression, respectively, in the GDA TCGA Pancreatic cancer cohort (n=176, log-rank test, *p<0.05, UCSC Xena).

## Discussion

Together, the presented data provide evidence that efficient cellular cholesterol removal mediated by small, lipid-poor reconstituted HDL particles reduce pancreatic cancer cell proliferation and growth and might hold the potential to attenuate the development and spread of the disease.

Although SR-B1 is highly expressed in the applied pancreatic cancer cell lines and significantly contributes to HDL-mediated cholesterol flux, forced efflux via ABCA1 increases the anti-tumor activity of those particles (Figure 7). In contrast, lipid-laden spherical HDL particles, which exhibit a higher affinity for SR-B1-directed lipid exchange at the plasma membrane, show reduced or insignificant ability to counteract cancer cell proliferation and tumor growth (Figure 7). Unidirectional, ABCA1-driven cholesterol efflux via small rHDL particles might thereby cause depletion of cellular cholesterol pools, eventually leading to decreased proliferation and viability of cancer cells. These results indicate that the HDL particle composition and thereby HDL functional metrics might determine its anti-tumor capacity.

**Figure 7.**
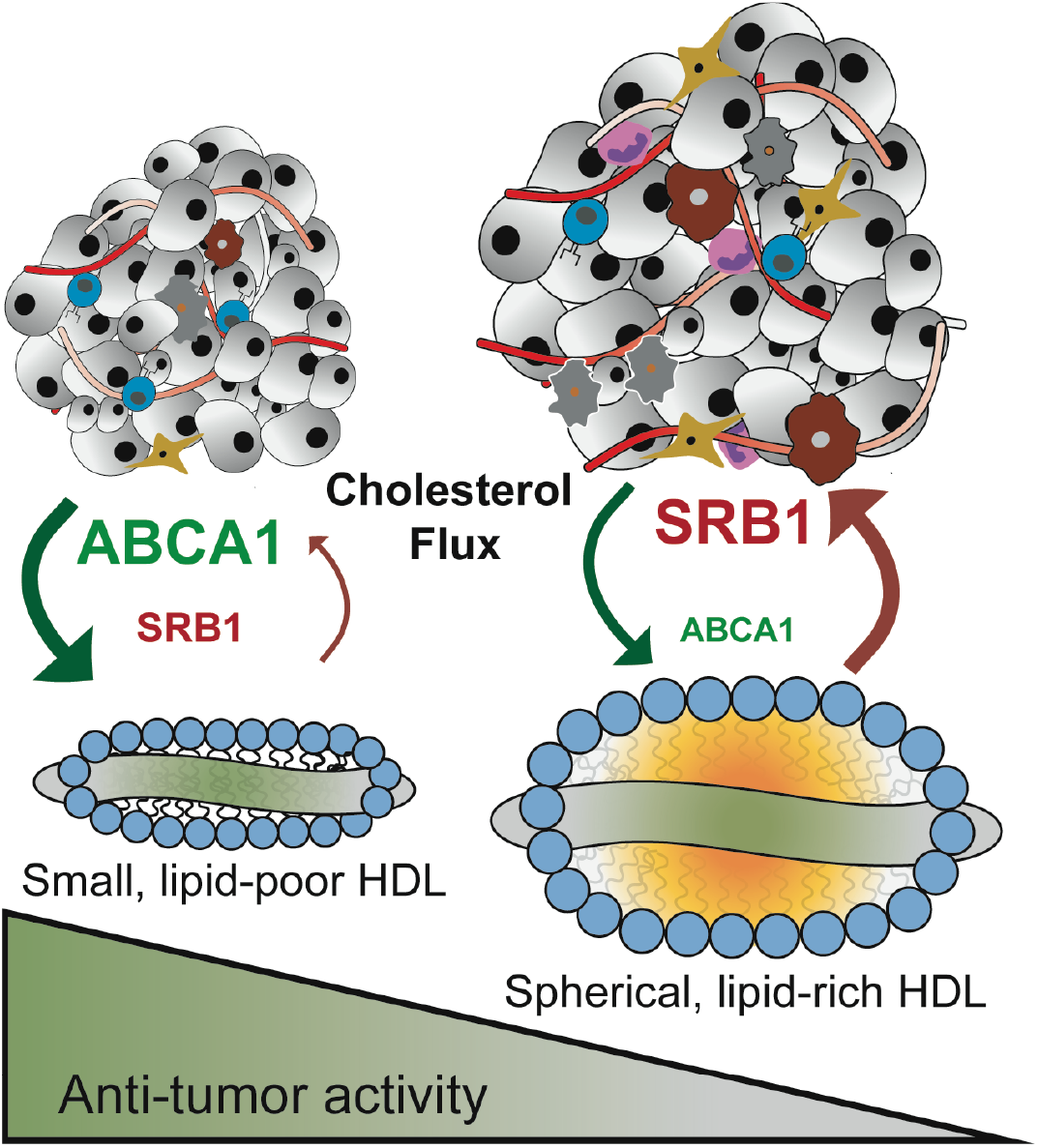
Small, lipid-poor HDL particles exert profound anti-tumor properties in PDAC models. Discoidal, lipid-poor reconstituted HDL, by engaging ABCA1-mediated net cholesterol efflux, reduce the viability and induce apoptosis of pancreatic cancer cells. In contrast, lipid-rich spherical HDL particles exhibit a higher affinity towards SR-B1, paralleled by a marked decrease in their ability to reduce cancer cell viability and inducing cancer cell apoptosis. Therefore, the particle-specific anti-tumor activity might be regulated by varying affinities of HDL subspecies towards SR-B1 and ABCA1, which are the dominant receptors expressed on PDAC cells that drive cholesterol flux at the plasma membrane. Increasing ABCA1-centered cholesterol depletion via discoidal, lipid-poor reconstituted HDL particles might be considered a valuable supplementary strategy to increase prognosis of PDAC patients in the future.

In the field of cardiovascular research, HDL functionality is currently under intense investigation, as the gold-standard plasma parameter, level of HDL-C, has been shown not to correlate with cardiovascular risk in interventional trials aiming to increase circulating HDL-C (39). One potential explanation for this discrepancy is the vast heterogeneity of the HDL particle pool. HDLs appear in the plasma as small, lipid-poor pre-β HDL and lipid-enriched ɑ-HDL particles. Thereby, the pre-ß fraction (mimicked by HDL1 particles used in this study), which only comprises about 5% of totals HDLs in the circulation, performs net cholesterol efflux from peripheral cells, predominantly macrophage foam cells, to eventually become ɑ-HDLs (9). This ɑ-HDL fraction is enriched in phospholipids, CE, and triglycerides, and acts as high affinity ligand for SR-B1, which, under physiological conditions, serves as receptor on hepatocytes that binds and sequesters HDL-associated cholesterol for excretion (31). SR-B1 in turn was previously shown to be overexpressed in many cancer entities including pancreatic cancer (40,41). Therefore, and as the results from this study indicate, ɑ-HDL particles might serve as cholesterol and CE source for cancer cells, eventually utilized as cellular fuel to drive cancer cell proliferation.

Efforts to artificially increase the cholesterol removing pre-β HDL fraction to drive ABCA1-mediated net cholesterol depletion from cancer cells might therefore provide a molecular axis that offers therapeutic potential for the treatment of PDAC. In support of this hypothesis, LXR agonists, which are potent activators of ABCA1 expression and thereby cholesterol efflux, reduced proliferation, cell cycle progression and colony formation of human PDAC cell lines (42). Moreover, LXR agonists have also been demonstrated to increase the expression of ABCA1 and the induction of the LDLR-degrading ubiquitin E3 Ligase IDOL, which leads to tumor cell apoptosis and a reduction in tumor growth in glioblastoma xenograft models (43).

Of note, high levels of LDLR and low levels of IDOL (*Mylip*) expression correlate with a worse prognosis in patients suffering from pancreatic cancer and other tumor entities (Figure 6 and Supplementary Figure 4). Cholesterol depletion by the use of statins was also shown to inhibit gallbladder cancer cell proliferation and sensitized those cells to cisplatin treatment, possibly by the inhibition of the DNA repair machinery (44). In view of those data, experiments which combine the administration of LXR agonists and efficient cholesterol acceptors such as pre-β-like rHDL particles with standard of care chemotherapy will provide valuable insights concerning the therapeutic applicability of the here presented preclinical findings.

As mentioned in the introduction, clinical studies indicate an inverse association of HDL-C with cancer incidence of multiple entities. Interestingly, data presented here point towards a concomitant decrease in the cholesterol efflux capacity of plasma samples from cancer patients (Figure 6). In addition to a reduction in HDL quantity, tumors might also be capable of influencing HDL functionality. HDL associated proteins such paraoxonase 1 (PON1) and serum amyloid A (SAA) as well as biochemical modifications of HDL structural components such as myeloperoxidase (MPO)-mediated nitration or chlorination of APOA1 are currently known to influence HDL’s reverse cholesterol transport capacity. The overexpression of PON1, an HDL-associated enzyme with potent anti-oxidative activity, has been shown to increase HDL-C efflux *in vitro* and reverse cholesterol transport *in vivo* (45). Of note, PON1 serum activity is reduced in cancer patients of various entities (40). SAA1 is an acute phase protein and transported predominantly on HDL in the bloodstream. Upon infection, SAA1 levels increase dramatically and its association with HDL has been demonstrated to reduce the lipoproteins’ anti-inflammatory properties and its cholesterol efflux capacity (40,46). Interestingly, certain cancer cell lines, tumor associated macrophages and pancreatic cancer-associated adipocytes produce large amounts of SAA (40,47). SAA levels were furthermore shown to directly correlate with disease progression, reduced survival rate and poor overall prognosis (40). In addition, macrophages and myeloid-derived suppressor cells accumulating in cancer patients express high levels of the enzyme MPO, a candidate enzyme that oxidatively modifies HDL, thereby reducing its cholesterol efflux capacity (39,48). Interestingly, MPO-mediated HDL modifications enhanced association of HDL with macrophages in cell culture and increased cholesteryl-ether transfer into target cells in an SR-B1-dependent manner (48). If this scenario is likely to happen in the tumor microenvironment, oxidatively modified HDL particles, while losing their cholesterol removal capacity, might serve as efficient cholesterol donors in SR-B1-expressing cancer entities such as pancreatic cancer.

In summary, the presented data demonstrate a potentially important role of HDL-mediated cholesterol efflux in reducing the proliferative as well as tumor initiating capacity of PDAC cells. Thereby, the HDL particle composition dictates its anti-tumor activity by regulating directionality of net cholesterol flow between cancer cells and the lipoprotein particle. Future studies, integrating the manipulation of cancer cell-specific HDL receptor expression as well as the detailed analysis of HDL particle functionality should be addressed to eventually implement cholesterol depletion as a combinatorial and supportive treatment modality in pancreatic cancer.

## Acknowledgements

The authors would like to thank Agnes Hunger, Franziska Pupp, Victoria Guggenberger and Anna-Lena Höbler for excellent technical assistance and Ernst Steyrer and Wolfgang Sattler (both Meduni Graz) for support and fruitful discussions in the process of conceptualizing and preparing this manuscript. RB was supported by an Erwin-Schrödinger fellowship of the Austrian Science Fund (FWF, J3664-B19). FU received a Werner Otto fellowship from the Werner Otto foundation. MW was supported by the Medical Faculty of the University of Hamburg (FFM program). SL is supported by the by the European Research Council (ERC) under the European Union’s Horizon 2020 research and innovation programme (Grant Agreement No. 758713) and by the Hector Foundation II.

## Notes

**Authors report no potential conflicts of interest.**

